# Rapid integration of face detection and task set in visually guided reaching

**DOI:** 10.1101/2023.12.07.570480

**Authors:** David Y. Mekhaiel, Melvyn A. Goodale, Brian D. Corneil

## Abstract

The superior colliculus (SC) has been increasingly implicated in the rapid processing of evolutionarily relevant visual stimuli like faces, but the behavioural relevance of such processing is not clear. The SC has also been implicated in the generation of upper-limb Express Visuomotor Responses (EVRs) on upper limb muscles, which are very short-latency (within ∼80 ms) bursts of muscle activity time-locked to visual target presentation. This reasoning led us to investigate the influence of faces on EVRs.

We recorded upper limb muscle activity from young healthy participants as they reached toward left or right targets in the presence of a distractor stimulus presented on the opposite side. Across blocks of trials, we varied the instruction as to which stimulus served as the target or distractor. Doing so allowed us to assess the impact of instruction on muscle recruitment by examining trials when the exact same stimuli required a reach to either the left or right. We found that EVRs were uniquely modulated in tasks involving face selection, promoting reaches toward or away from faces depending on instruction. Follow-up experiments confirmed that this phenomenon required highly salient repeated faces, and was not observed to non-facial salient stimuli nor to faces expressing different affect. We conclude that our results attest to an integration of top-down task set and bottom-up feature detection to promote rapid motor responses to faces at latencies that match or precede the arrival of face information in human cortex.

**STATEMENT OF SIGNIFICANCE:** The tecto-reticulo-spinal pathway is hypothesized to mediate the express visuomotor response (EVR). This study extends this hypothesis by demonstrating that face detection in the subcortex impacts low-latency movement via the EVR at latencies preceding cortical activity for face perception. To date, this constitutes the most direct evidence for direct behavioural relevance of rapid face detection in the brainstem. Further, we find that this response can be modulated by task context, allowing for different instruction-based responses given the exact same visual stimulus and implicating top-down cortical control of the EVR.

## INTRODUCTION

Orienting to stimuli is so fundamental to an organism’s survival that it has largely driven the evolution of our senses. When time is of the essence, it is thought that the superior colliculus (SC) can rapidly transform visual inputs into motor outputs to generate orienting reflexes by coordinating the rapid recruitment of muscles to facilitate low-latency gaze shifts (Corneil & Munoz, 2014; Gandhi & Katnani, 2011). A growing body of evidence suggests that the SC, via projections of the tecto-reticulo-spinal network (TRSN), can also influence limb muscle recruitment. In particular, it is thought that the SC and TRSN may mediate a reflex known as the express visuomotor response (EVR) in the muscles of the upper limbs that is temporally linked to the presentation of a visual stimulus. The EVR is short in latency (80-120ms) and can be expressed as either an increase or decrease in muscle activation, depending on the location of the stimulus (Pruszynski et al., 2010). By modifying limb muscle activity, the EVR contributes to the initiation of shorter latency reaches to suddenly appearing stimuli (Gu et al., 2016; Kozak et al., 2019).

Additional support for the hypothesis that EVRs are mediated by the SC and TRSN comes from the fact that low-spatial frequency and high-contrast stimuli not only engender larger responses in movement-related layers of the SC and more frequent express saccades (Chen et al., 2018; Marino et al., 2012), but also evoke larger EVRs in the upper limbs (Kozak et al., 2019; Kozak & Corneil, 2021). In addition, priming and task set can affect the production of both express saccades (Meeter & Van der Stigchel, 2013) and EVRs (Contemori et al., 2021, 2023).

Recent work has expanded the understanding of the brainstem feature-detection system by showing that evolutionarily-relevant stimuli, like faces and snakes, evoke shorter latency orientation movements than other kinds of visual stimuli (Almeida et al., 2015; Bannerman et al., 2009, 2010; Martin et al., 2018). In addition, neurons in the primate SC and pulvinar exhibit lower latencies (∼50ms) and larger bursts in response to faces than other visual stimuli (Almeida et al., 2015; Nguyen et al., 2013, 2014, 2016; Yu et al., 2023). These subcortical networks are thought to play a crucial role in the detection of faces and facial expressions in patients with lesions of the primary visual cortex (Celeghin et al., 2019). Taken together, these observations suggest that the SC responds to faces independently and well before face areas in the cerebral cortex (Bentin et al., 1996; Collins & Olson, 2014; Decramer et al., 2021; Kato et al., 2004). Consistent with the fact that SC detects faces faster than other visual stimuli, faces evoke shorter-latency and more frequent express saccades (Salvia et al., 2020; VanRullen & Thorpe, 2001). Curiously, the saccadic preference for faces is only detectable within saccadic choice paradigms, wherein participants look to an instructed target presented synchronously with a distractor (Crouzet & Thorpe, 2011).

If EVRs in the upper limbs are also mediated by the SC and TRSN, then one might expect that they too will be affected by (1) the presentation of specific targets, such as faces, and (2) task set. In short, we predict Larger EVRs in the muscles of the arm in reaching tasks featuring faces, resulting in more vigorous muscle recruitment and thus more rapid reaching movements.

Our results support our predictions, supporting the conjecture that face detectors in the SC/TRSN mediate the initiation of EVRs when face targets are presented. Further, and most intriguingly, top-down control, based on instruction or task set, can modify how the EVR directs movement towards or away from faces.

## METHODS

### Participants

Five experiments were conducted in this study. University students were recruited using the Psychology Research Participant Pool. All participants provided informed written consent and were free to withdraw from the study at any time. Participants were compensated financially or with research credits required for course completion. All procedures were approved by the Health and Science Research Ethics Board at the University of Western Ontario. All participants had normal or corrected to normal vision and reported no neurological or motor disorders.

For experiment 1, data from a cohort of 20 participants were collected (female: 8, male: 12; mean age: 23yr, SD: 4.68). Experiment 2 and 3 were completed in the same session by a second cohort of 20 participants (female: 9, male: 11; mean age: 18.3yr, SD: .86). Finally, experiments 4 and 5 were completed in the same session by a third cohort of 20 participants (female: 9, male: 11; mean age: 18.45yr, SD: .76). No participant participated in more than one of the aforementioned three cohorts. These cohorts do not include data from 6 participants who were excluded: 1 for a resting tremor, 1 for failing to understand the task, and 4 for having a trial exclusion rate above 50%.

### Apparatus

The experiments were conducted using a Kinarm Endpoint Robot (Fig. 1A; BKIN technologies, Kingston, ON, Canada). A Propixx projector (VPixx, Saint-Bruno, QC, Canada) integrated within the Kinarm platform enabled presentation of high-quality visual stimuli and ensured reliable event timing. A photodiode was employed to verify the exact time the stimuli appeared in each trial. Behavioural tasks were generated using Stateflow and Simulink within MATLAB (version R2021a, MathWorks Inc., Natick, Massachusetts, United States of America). The experimental paradigms were projected onto a horizontal surface which was approximately at shoulder level to the participants when seated.

**Figure 1.**
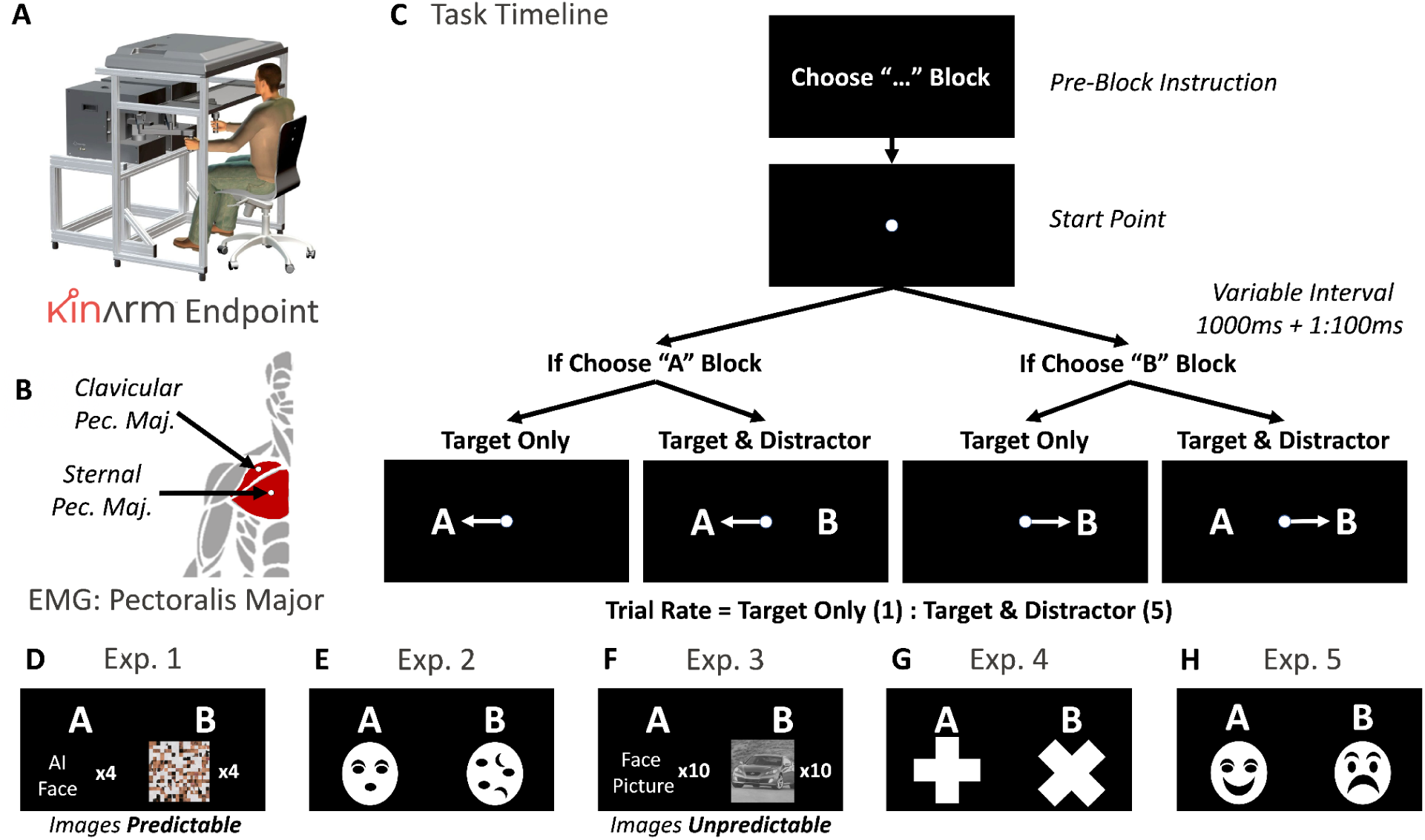
Experimental apparatus & task. (A) All data was collected while participants made planar reaches with their right arm in a Kinarm End-Point Lab. (B) Surface EMG activity was recorded from the sternal and clavicular heads of right pectoralis major muscle, which increases/decreases in activity prior to leftward/rightward reaches. (C) Participants were instructed to reach towards one of two targets (or target categories) at the beginning of each block. Participants initiated each trial by moving the Kinarm manipulandum to the start point. After a variable interval of time the target appeared to the right or left either on its own (Target Only; ∼17% of all trials) or with a distractor on the opposite side (Target & Distractor; 83% of all trials). (D-H) stimuli employed in each of the five experiments. Across blocks, each image or image category could serve as either the target or the distractor. In experiment 1 and 3, image categories were used (e.g., different faces could appear in conditions involving faces). In experiment 1, the target & distractor images changed *between blocks*, but remained predictable *within* a block of trials. In experiment 3, the target and distractor images were randomized *within* a block of trials, and hence were unpredictable. For the purpose of BioRxiv submission, natural-appearing face stimuli have been replaced with text throughout the manuscript.

All participants made reaches with their right arm, regardless of handedness. Surface electromyography (EMG; Delsys Inc. Bagnoli-8 system, Boston, Massachusetts, United States of America) recordings of the right pectoralis major muscle were acquired in two locations: the clavicular head and the sternal head of the muscle (Fig. 1B), which were identified using palpation. EMG data was filtered using a high and low pass filter of 20 and 450 Hz, respectively, and then digitized at 1000 Hz onto the Kinarm system.

Participants completed the experiments using a manipulandum under the display surface to control a cursor (Fig. 1C). An occluder obscured view of their hand and of the manipulandum. A constant force of 2N towards the participant and 5N towards the right was applied to the arm by the manipulandum throughout the experiments to induce baseline muscle activity in the right pectoralis. Kinematic data from the manipulandum was recorded at a rate of 1000Hz by the Kinarm system. Eye movement data was not recorded.

### Experimental design

Five experiments were designed to test how EVR activity differed in response to (1) different types of visual stimuli and (2) task set based on block instruction delivered to the participant. Since it is our hypothesis that the EVR is mediated by the SC, the experiments drew inspiration from studies investigating saccadic eye movements and the visual response properties in the SC.

All experiments required the participant to make reaching movements towards a prescribed target or target category (Fig. 1D-H). Participants initiated each trial by aligning the Kinarm manipulandum within a 2cm diameter starting point for 500ms. They were instructed to fixate their gaze on the start point until the reach target appeared. Following a variable time of 500-600 ms (hence the total time at the start point was between 1000 to 1100 ms), the target appeared either to the left or to the right of the starting point. The temporal uncertainty was used to reduce the occurrence of anticipatory reaches. In 17% of trials, the target appeared on its own. In the remaining 83% of trials,a target and distractor appeared simultaneously. The ratio of target only trials to target & distractor trials was 1:5. The center of the target and/or distractor was 10 cm away from the center of the starting point, this translated to ∼9.5 degrees of visual angle from the center of the starting point to the center of the target or distractor. Participants then responded. The trial ended once the cursor contacted the target or the distractor.

Before each experiment participants completed a practice session of 100 trials. Each experiment then consisted of 480 trials including 400 target & distractor trials and 80 target only target trials. The experiments were broken up into eight blocks of 60 trials (50 target & distractor, 10 target only), after which the instructed target was alternated (e.g. a “choose face” block was followed by a “choose car” block). Following four experimental blocks, the participants were given a timed 5-minute break. Within a given experiment, block order was alternated between participants. In experimental sessions where participants completed more than one experiment, the order of the experiments was likewise alternated between participants.

### Experimental stimuli

For experiment 1, we examined the EVR to natural looking images versus their scrambled counterparts. We used “Generated Photos” AI to generate faces of four individuals, 2 men and 2 women of varying ethnic backgrounds. To generate the scrambled counterpart, each image was broken up into a grid and the squares within were randomized. Within each block only one of the four faces with its corresponding scrambled image were presented so that the target was predictable to the participant.

In experiment 2, we used a simplified version of experiment 1, using geometric shapes arranged to either resemble a face, or arranged randomly into an abstract figure. The use of geometric shapes in experiment 2 allows us to control for spatial frequency and luminance without reduction in image clarity. High contrast images of this type have been found to elicit higher neural activity in the SC of the monkey when compared to natural images of faces (Nguyen et al., 2013, 2014, 2016; Van Le et al., 2020).

In experiment 3, we selected images from the same visual database (Corel Photo Library) used by Thorpe and colleagues to investigate saccades (Crouzet et al., 2010; Crouzet & Thorpe, 2011). Images in this database were normalized for variations in luminance and spatial frequency. Ten images of faces and ten images of cars were selected. The images of faces selected included 5 men and 5 women of varying age and ethnicity. The pictures of faces and cars were made unpredictable through randomization, with each image appearing 1 or 2 times within a single experimental block either as a target or distractor. In alternating blocks, participants were instructed to reach either towards the face or the car.

In experiment 4, we investigated if the EVR differentiates between the same geometric shape in two different orientations (e.g., a “+” or “x.”). In experiment 5, we investigated if the EVR is influenced by geometric faces displaying *happy* or *sad* affects.

### Kinematic analysis

All analyses were completed using MATLAB (version R2021a, MathWorks Inc., Natick, Massachusetts, United States of America). On each trial, a white stimulus that was unseen by the participant appeared simultaneously with target and distractor onset and was detected by a diode. All kinematic and EMG data was aligned to diode onset.

Inclusion and exclusion criteria for trials was determined using the kinematic output from the Kinarm manipulandum. The second derivative of the horizontal position within the workspace was used to determine the horizontal acceleration of the participant’s arm. The mean acceleration for the 100ms preceding target onset to 50ms following target onset for all trials was used to identify the baseline variation in acceleration for each participant. Movement onset was detected when hand acceleration increased beyond the 95% confidence interval of the baseline mean acceleration for the participant. A graphical user interface was used to confirm and adjust, if necessary, the onset and offset of movements.

Trials were labelled as “false start” and excluded if the participant initiated a movement from 100ms preceding target onset to 120ms following target onset. We excluded “too fast” trials where the reaction time was between 120ms to 150ms. Trials were labelled as “wrong way” and excluded if at any point following target onset the participant’s hand moved towards the distractor. These two criteria ensured that EMG activity during the EVR epoch was not contaminated by voluntary movement related activity. Exclusion of “too fast” trials did not have any impact on the findings of the study, since the mean reaction time across all experiments for target & distractor trials was greater than 200 ms.

An additional spatial analysis was conducted to ensure that muscle recruitment related to anticipation did not influence our analysis of the EVR. Since a participant’s hand may drift outside of the starting position slowly, thus remaining below the acceleration threshold, their hand may leave the starting position without the preceding analysis registering a movement. Given that both the cursor and starting position had a radius of 1 cm, we also identified trials where the center of the cursor was 2 cm or more from the middle of the starting point before the participant made their final movement towards the target. Such trials were labelled as “unstable” and were also excluded. The rate of “unstable” trials exceeded 50% for 5 participants in experiment 1, 6 participants in experiments 2 and 3, and 2 participants in experiments 4 and 5. For these participants, we opted to retain the data from these subjects including the unstable trials. However, we re-ran all analyses after excluding all data from these participants, and confirmed that doing so did not alter our main findings (results not shown).

### EMG filtering and normalization

Offline, EMG data was full-wave rectified and then filtered by a 7-point moving average filter. EMG data was normalized on a trial-by-trial basis to the EMG activity counteracting the constant torque applied to the arm by dividing the EMG signal over the entire trial by the mean activity in the 100ms preceding the stimulus onset. Since results did not vary between the sternal and clavicular recordings, the two EMG recordings were then averaged.

### Analysis of the EVR

Previous work analyzing the EVR typically contrasts trials where a stimulus appeared to the right or left. In the current study, we are aiming to determine whether the EVR on target-distractor trials is influenced by the features of the presented stimuli and by the block instruction to reach towards a specific stimulus. Thus, a key comparison will be to examine EMG activity when the exact same stimuli are presented to the retina: we predict that the EVR on the pectoralis muscles should be larger when the block instruction requires that the participant reach to the stimulus on the left rather than the right (recall that the right pectoralis muscles are recruited for leftward, cross-body reaches).

Based on previous behavioural and neurophysiological studies, we expect that the influence of stimulus features and block instruction on the magnitude of the EVR to be relatively small within any given subject. We therefore used group analysis methods which are more sensitive to small differences, relying on paired comparisons using both frequentist and Bayesian approaches.

For our frequentist approach, we conducted a timewise paired T-test to analyse the effect of block instruction on mean muscle activity for each millisecond within the EVR epoch (80-120 ms following stimulus onset). This was followed by a Benjamin and Hochberg false discovery rate correction (p-crit=0.05).

Bayesian statistics, although having a limited history of implementation in the literature within similar contexts, have many advantages. Unlike frequentist statistics with which we can only reject a null hypothesis or fail to reject it, Bayesian statistics allows us to quantify the level of certitude with which we believe a null hypothesis (M_0_) or alternative hypothesis (M_1_) is true. Additionally, sequential Bayesian statistics allow us to reiteratively sample participants until the “*Bayes factor,*” (BF) or the ratio between the two-competing hypotheses, remains stable within a given value range. Providing such stability is reached, the final statistical conclusion would not change with the addition or removal of a small number of participants. The visualization of Bayesian statistics is also a good test to see if a given result is consistent with iterative sampling. Finally, Bayesian statistics are non-parametric and therefore are a more flexible model for data analysis.

Bayesian analyses were conducted in Matlab R2021a using the function BayesFactor v.2.3.0 (Krekelberg, 2022). Our Bayesian analysis consisted of a one-sided paired T-test to compare the effects of block instruction on EMG magnitude. The model uses a multivariate Cauchy prior with a scale of 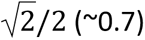 to calculate the JZS-BF, as described by (Jeffreys, 1998; Zellner & Siow, 1979). The location of the instructed target was the independent variable and the EMG magnitude was the dependent variable. Given EMG data for reaches to the left (*EMG*_*Left*_) and reaches to the right (*EMG*_*Right*_), the BF was calculated using two different models (our two hypotheses):

***Null hypothesis*** ( *M*_0_) :

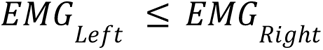

***Alternative hypothesis*** (*M*_1_):

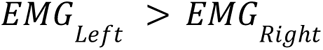

Given data (*D*), the Bayes factor (*BF*) can be calculated as such using the posterior probabilities (*Pr*) of both models. For the purpose of this study, we calculated the inverse of the standard *B*_01_(*B*_10_), meaning that the BF is directly proportional to the probability of M_1_.

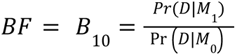

This analysis was run sequentially with the addition of each participant (from N=1 to N=20) for a time spanning from 500 ms before to 1000 ms after target onset. The validity of the final BF for each time point (including all participants) can be verified by looking at how the BF changed over time with the iterative inclusion of each participant.

To demonstrate this group bayesian analyses, we present an analysis using the target only conditions for experiment 1 (Fig. 2). The left most column (Fig. 2A) shows the mean EMG for correct target-only face trials where the target was on the left or right for the first two participants, in the order of sampling. As previously discussed, one advantage of bayesian analyses is that they can be run sequentially, therefore allowing us to visualize how the BF changes with iterative sampling and across each time point. The second column demonstrates the BF across time with the inclusion of the first participant only (N=1) and the first two participants (N=2). The boundaries in the y-axis describe the heuristic classification scheme for the BF indicating level of support for either *M_0_* or *M_1_*, from “anecdotal” to “extreme” (Jeffreys, 1998). These values are also visualized in the heatmap (Fig. 2B), where the EVR epoch (80-120ms) is bolded. The heatmap is scaled for a BF of 0 to 10 (blue to red), meaning that red value would indicate, at minimum, “strong” support for our alternative hypothesis.

**Figure 2.**
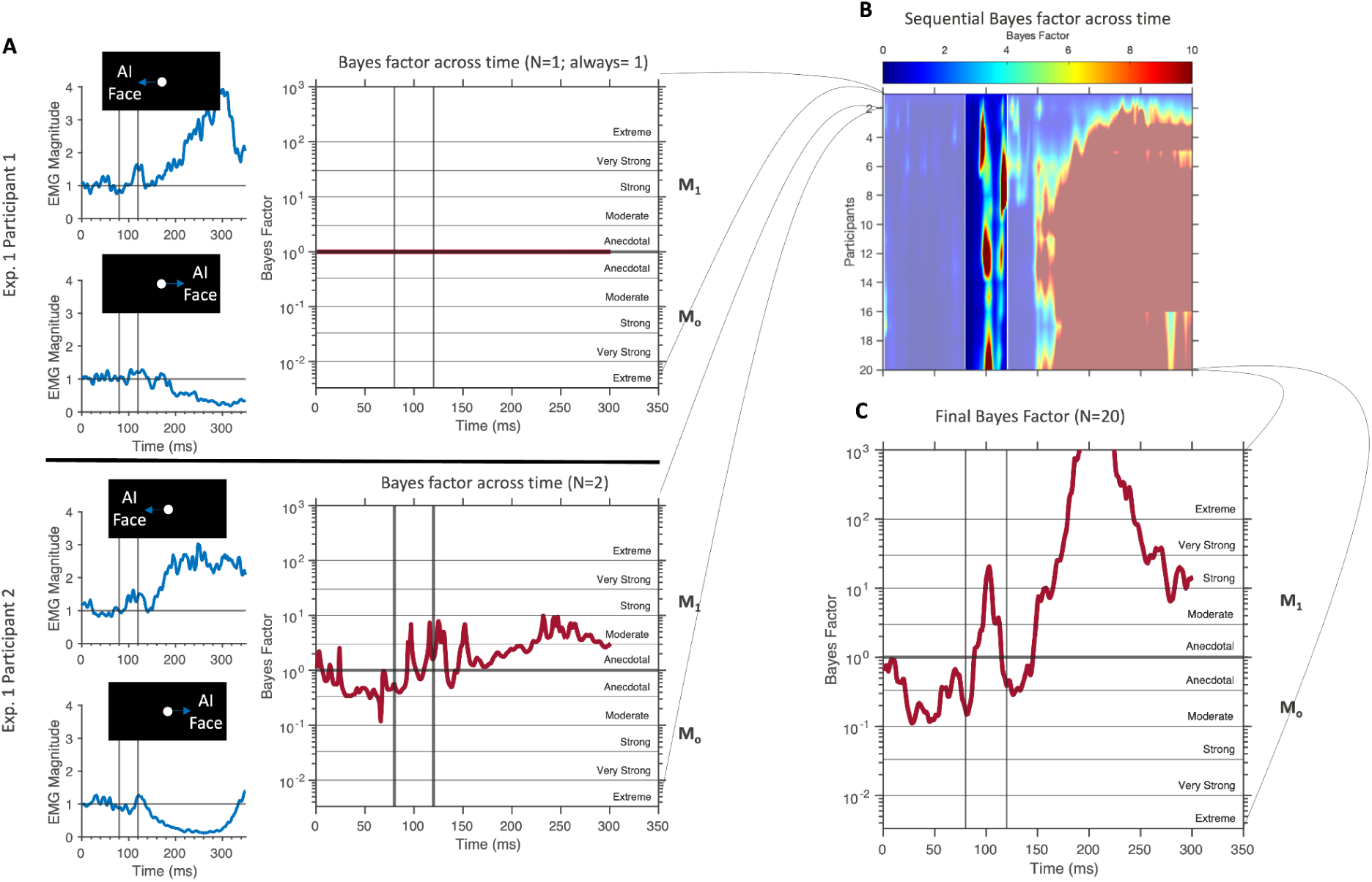
Demonstration of sequential Bayesian analysis of EMG recruitment across time using target-only trials in Experiment 1. (A) Blue traces in the left-most column show mean EMG activity for two participants for target-only trials to the left (upper subplots) or right (lower subplots). The red lines in the second column from the left show the Bayes factor through time. When data from only one subject is considered (upper plot), the Bayes factor is always equal to 1. Given the first participant’s data as a posterior, with the inclusion of the second participant, the BF indicates a moderate effect of the conditions on the EMG activity during the EVR epoch, favouring *M1* as expected since the right pectoralis major muscle is recruited for leftward reaches. (B) Such time courses are created for the sequential inclusion of each participant, which can be viewed by a heatmap where the colour displays the value of the Bayes factor as a function of time after stimulus presentation (x-axis) and the number of included participants (y-axis). A value of 10 (10^1^) or higher indicates a “strong” impact of target location; note how evidence for this occurs during the EVR epoch (80-120 ms), as indicated by the red banding in the bolded region. (C) The Bayes factor as a function of time when data from all 20 participants is included. Here, a Bayes factor between 10 and 30 during the EVR epoch indicates a “strong” likelihood of the alternative hypothesis; that muscle activity in the right pectoralis major is higher when a face is presented to the left vs to the right. The Bayes factor gets even higher (> 100, or 10^2^) during subsequent phases of muscle recruitment (> 150 ms after stimulus onset), which is expected given the recruitment of right pectoralis major for leftward reaches.

To ensure the validity of the final BF we should observe consistent values with sequential sampling (the addition of new participants). For instance, this is confirmed through the consistent red banding during the EVR epoch. Plotting the final BF across time with the inclusion of all participants (N=20) demonstrates that we observe moderate evidence for *M_0_* before the EVR epoch. We then observe two BF peaks, the first during the EVR epoch indicating “strong” support for *M_1_*and the second during the voluntary response epoch (150ms+) indicating an “extreme” likelihood of *M_1_*. The distinction between the two BF peaks is consistent with the difference observed during the EVR epoch not being driven by the voluntary motor response. These results are in line with our subjective observations of EMG activity during target only trials (Fig. 2A), where there are distinct increases of activity during leftward reaches and decreases in activity during rightward reaches in both the EVR and voluntary movement epochs.

## RESULTS

Recordings in the primate SC show that presentation of face or face-like stimuli can influence the vigor of visual responses within <50ms (Almeida et al., 2015; Nguyen et al., 2014, 2016; Yu et al., 2023). Given that the tecto-reticulo-spinal pathway is hypothesized to mediate the EVR on the upper limb (Corneil & Munoz, 2014; Pruszynski et al., 2010), we investigated if presentation of face or face-like stimuli influence the magnitude of the EVR. We recorded surface EMG from the right pectoralis major muscle, which is active during leftward movements, during a reaching choice task. Across five experiments human participants were instructed to reach toward a target defined by various features, often in the presence of the distractor presented on the opposite side. In different blocks, we interchanged the instructed target, allowing us to compare the influence of block instructions on the EVR given the exact same visual inputs.

### Consistent error rate across experiments

Across all five experiments, ∼75% of target & distractor trials met our acceptance criteria (Fig. 3A). Of the trials that did not meet our acceptance criteria, we rejected ∼2% of all trials as anticipatory, ∼1% of all trials that moved all the way to distractor, and ∼22% of trials composed of multiple movement segments (e.g., the subject initially moved to the distractor and then moved to the target). Rejecting these trials ensured that participants consolidated the block instruction to move toward a specific target feature and ensured that EMG activity related to anticipation did not influence muscle recruitment during the EVR window (80-120 ms after target onset). As will be discussed at the end of the results section, inclusion of 22% of trials where the participant first first inappropriately moved to the distractor did not change the overall interpretation of our results.

**Figure 3.**
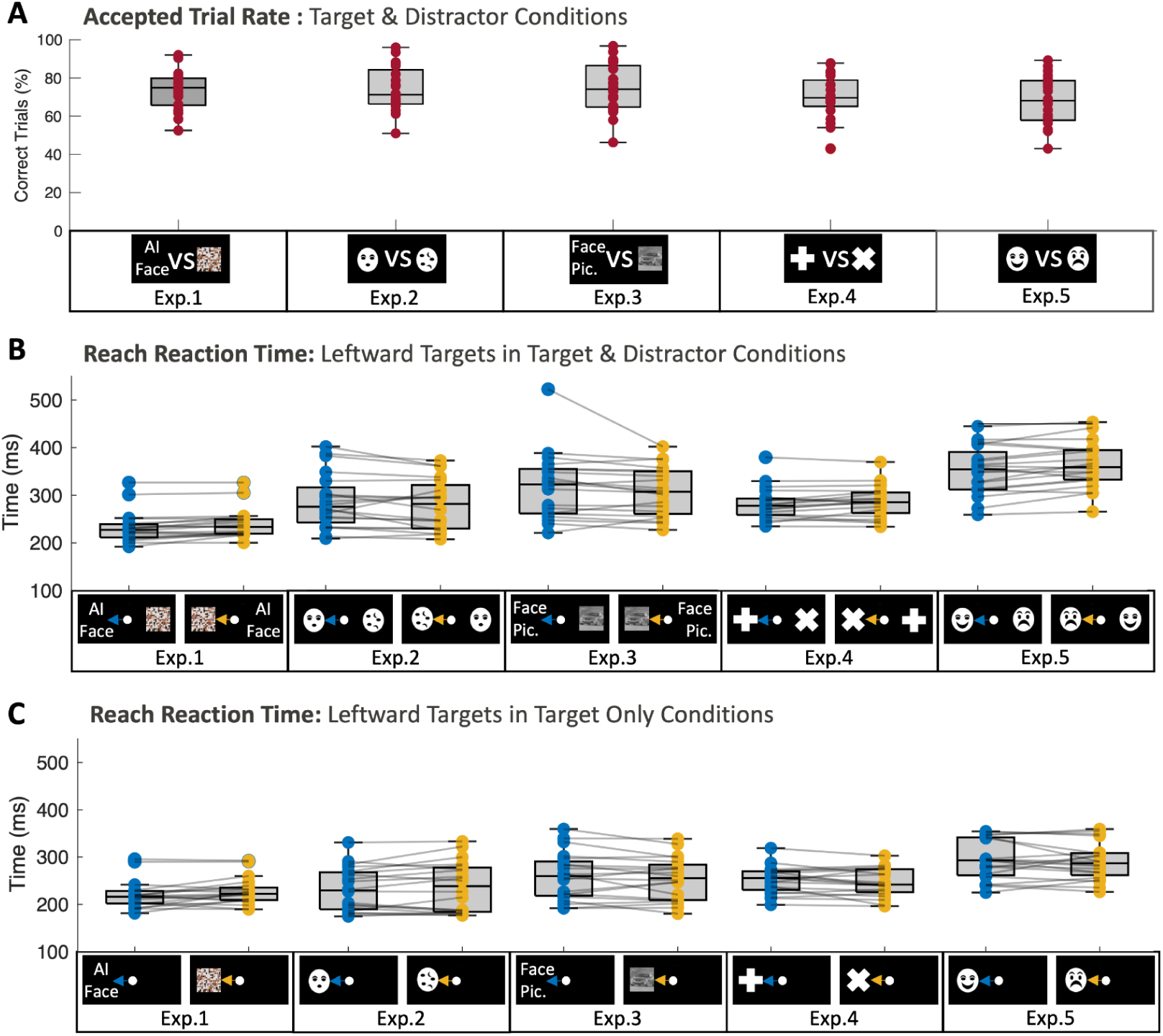
Analysis of performance. (A) Error rate for target-distractor conditions for experiments 1-5. B, C. Mean leftward reaction times for correct movements in each condition for each experiment for either target-distractor (B) or target-only (C) trials. Since the manipulandum was loaded to the right, we analyzed leftward reaches. We used repeated measures ANOVA to analyse the effects of target types and presence of a distractor on RT and error rate. Each dot and line connects data from a single participant.

We performed a two-way repeated measures ANOVA in each experiment to analyze the effect of target type and the presence of the distractor on error rate. The analysis showed that target type did not have a statistically significant effect on error rate in any experiment (all p >.458), while the presence of the distractor resulted in a statistically significant increase in error rate across all experiments (all p <.001). There was no statistically significant interaction in any experiment (all F(1, 76) < .9, p >.345).

### Slower reaction times in target & distractor conditions

Next, we examined the pattern of reaction times (RTs)across all experiments (Fig. 3B,C), focusing on leftward reaches since we measured the recruitment of the right pectoralis major muscle. Within each experiment, we are specifically interested in whether RTs vary as a function of the block instruction, and by the presence or absence of a distractor. We therefore ran a two way repeated measures ANOVAs to examine the effect of instructed target type and the presence or absence of the distractor.

Although we predicted that limb movements towards faces would be initiated more rapidly than towards the scrambled image, we did not observe any statistically significant effect of the instructed target on RT (all p >.288) in any experiment. The presence of the distractor did have a statistically significant effect on RT, delaying RTs in experiments 2-4 (all p <.001; this effect approaches significance in experiment 1 (p =0.06)). There was no statistically significant interaction in any of the experiments (all F(1, 76) <.44, p >.509).

### The EVR persists on target-distractor trials

Recall that participants performed the task while their right arm was loaded with a constant force that increases the recruitment of right pectoralis major. In figure 4A, we show the data from a representative participant in experiment 1. As a consequence of the loading force, the EVR on target-only trials is easily identified as a brief increase or decrease in EMG activity ∼100 ms after target presentation to the left or right, respectively. Even on single trials, this initial phase of muscle recruitment is clearly more aligned to target onset than movement onset. The representative example in Fig. 4A displays a prominent rebound of EMG activity following rightward target-only presentation, similar to what has been observed previously (Wood et al., 2015). The key features of the EVR observed on target only trials mirror those described previously (Contemori et al., 2021; Gu et al., 2016; Kearsley et al., 2022; Kozak et al., 2019, 2020; Kozak & Corneil, 2021; Pruszynski et al., 2010). This is followed by a phase of voluntary muscle activity >150ms after target onset, in which an increase or decrease in muscle activity of the right pectoralis directs the arm to the left or right, respectively.

**Figure 4.**
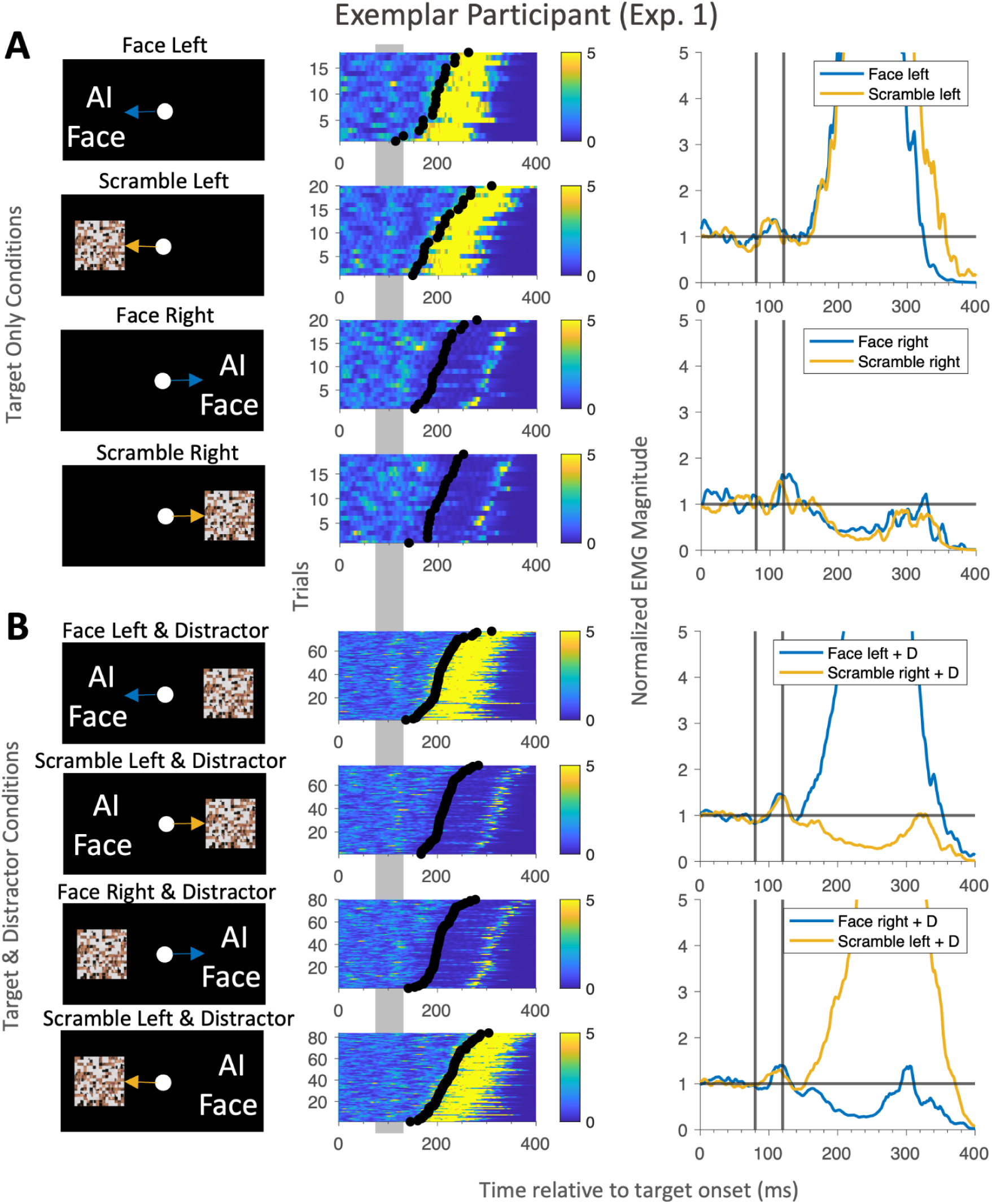
Exemplar participant in experiment 1. Representation of muscle recruitment for (A) target-only conditions and (B) 4 target & distractor conditions. Middle column shows data in a heatmap, where each row depicts single-trial normalized muscle recruitment in color aligned to target onset (0 ms), ordered by RT (black circles). The EVR epoch is shaded in grey. Rightmost column shows normalized muscle activity with the EVR epoch (80-120ms) demarcated by the vertical black lines.

Figure 4B shows data from conditions where the target & distractor were presented simultaneously on opposite sides. Somewhat surprisingly given the presentation of diametrically opposite stimuli, we still observed a prominent band of recruitment in the EVR epoch (80-120 ms) on target-distractor trials, which was also clearly separable from the subsequent phase of reach related activity. Importantly, this first phase of recruitment was not obviously lateralized and was largely similar across all four possible configurations of stimuli and block instruction. This observation resembles a recent report of EVRs following the presentation of identical diametrically opposed visual stimuli (Selen et al., 2023). Given that the timing of such activity and the general profile of recruitment (e.g. an ∼30 ms increase in activity more time-locked to stimulus than movement onset) is very similar to the EVR on target only trials (Fig. 4), this shows that the EVR persisted in this participant on target-distractor trials. Similar patterns were observed in all participants, in all experiments.

### Task set modulates the EVR in face related reaching tasks

Having established that the EVR persists in the target & distractor conditions, we next present an analysis of the effect of block instruction in these conditions. As shown in Fig. 1, our experimental structure, which was inspired by the 2011 study of Crouzet and Thorpe, enables us to compare the EVR given the exact same visual images but different block instructions. Consider for example trials in Experiment 1 where the ‘Face’ is presented on the left and the ‘Scramble’ is presented on the right; participants should reach leftward in the ‘Choose Face’ block but rightward on ‘Choose Scramble’ block. In the context of the EVR, comparing such trials isolates any effect of the block instruction. Our hypothesis predicts larger EVR on the right pectoralis major muscle when the face target is to the left versus the right.

#### Group analysis: Experiment 1

We start with the group outcomes from Experiment 1, which contrasted the EVR to an expected face versus a scrambled image of this face. For each participant, we quantified the magnitude of the EVR by integrating EMG activity in a window from 80-120ms (Fig. 5A), and then plotted the magnitude of the response to the exact same visual stimulus as a function of whether the instructed target was a face or not. As shown in the first two rows of Fig. 5A, we observed a significantly larger EVR across our sample to a leftward face in the ‘Choose Face’ block (i.e, when the participant subsequently reached to the left) than in the ‘Choose Scramble’ block (i.e., when the participant subsequently reached to the right; p<0.01). In contrast, a similar influence of block instruction was not seen when the scrambled image was present to the left (Fig. 5A), as we observed comparable magnitude EVRs in both the ‘Choose Face’ and ‘Choose Scramble’ blocks (p= .803).

**Figure 5.**
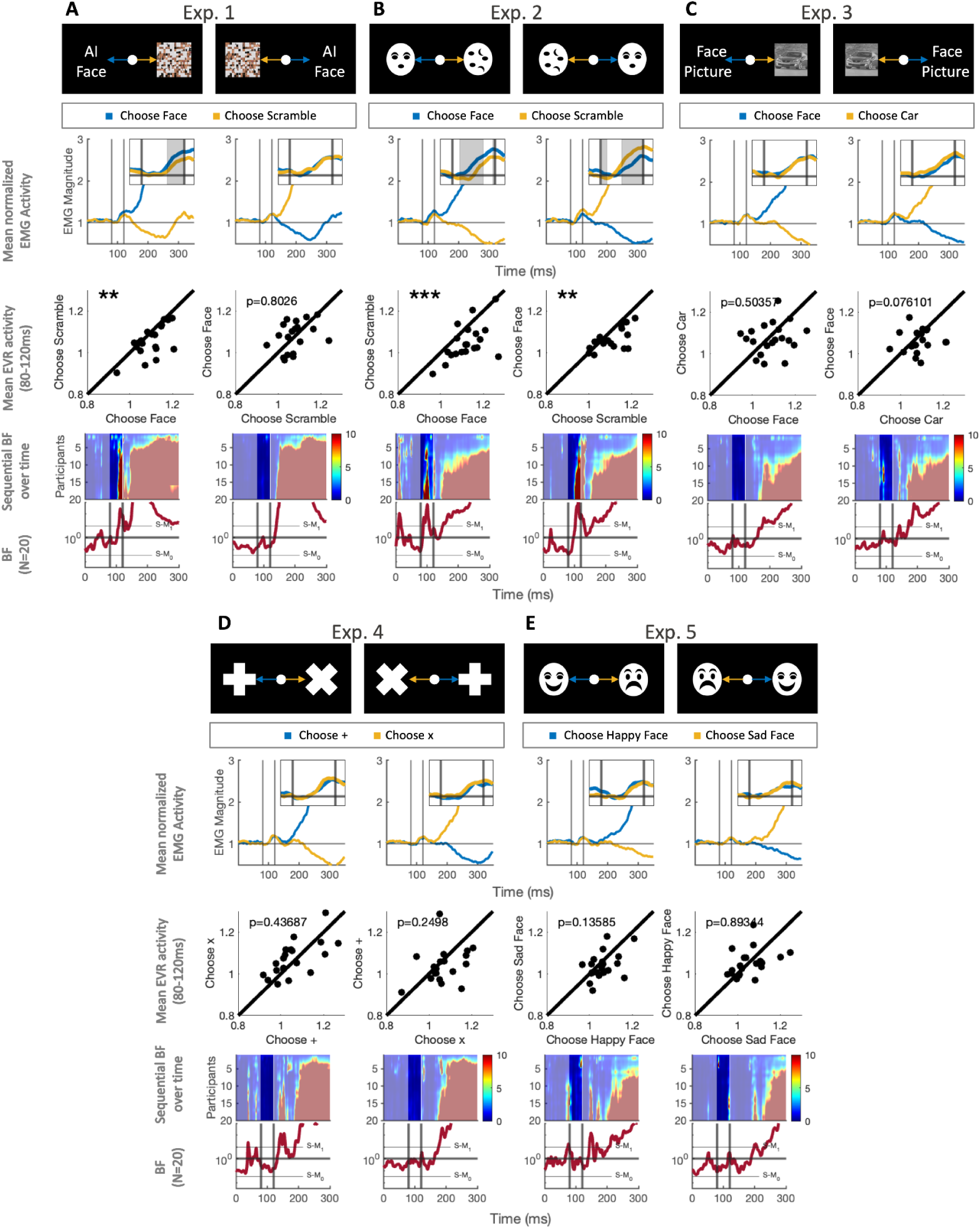
Group Frequentist and Bayesian Analyses for Experiments 1-5. *The first row* for each experiment indicates the mean of mean normalized EMG activity for all participants in target & distractor conditions. Each column contains statistical comparisons of conditions where the exact same images were presented to the participant, but where the participant had received different block instructions preceding the trial (e.g. reach to the face Vs. reach to the scramble in Experiment 1). The EVR epoch (80-120ms) is demarcated with 2 black lines. Grey shading in the EVR epoch signifies time points identified as significantly different, using a rolling paired two tailed t-test followed by a Benjamin Hochberg false discovery rate correction (p-crit=0.05). *The second row* shows a scatter plot of the mean muscle activity for each participant for each of the two conditions. Significant observations or p-values from a paired two-tailed t-test (*=p<0.05, **=p<0.01, ***=p<0.001). *The third row* for each experiment shows a BF for a one-tailed t-test to identify time points at which block instruction impacted EMG magnitude. An increase in the BF above 10^0^ (or 1) indicates higher likelihood of *M_1_*; i.e. that the EMG magnitude was higher when the participant was instructed to reach towards the target that is on the left vs. the target on the right. A BF lower than 10^0^ indicates a higher likelihood of the null hypothesis. The BF changes over time relative to stimulus onset and with the addition of every sequential sample (participant). Red banding during the EVR epoch, shows that the Bayes factor consistently indicates a “strong” likelihood of the M_1_. *The fourth row* for each experiment shows the BF across time with the inclusion of all participants (N=20). The three horizontal lines indicate BF= 10^0^ (no evidence for M_1_ or M_0_) and “strong” evidence for M_1_ (BF = 10/1) and M_0_ (BF =1/10) are indicated above and below 10^0^, respectively.

The third and fourth rows of Figure 5 show the BF factor for a paired t-test plotted across time with sequential sampling (upper subplot) and the final BF (N=20) across time (lower subplot). As reflected by the red banding during the EVR epoch in the upper plot, there was a consistent effect of block instruction with sequential sampling (the addition of new participants to the analysis). The final BF peaks during the EVR epoch at 113ms with a BF=272 indicating an “extreme” likelihood of the effect of block instruction on EVR expression when the face is presented to the left in the ‘Choose Face’ block. However, only “anecdotal” evidence of an effect was identified when the scrambled image was presented to the left in the ‘Choose Scramble’ block (BF= 1.17, t=98ms).

#### Group analysis: Experiment 2

Serving as a follow-up to experiment 1, experiment 2 employed high-contrast face symbols, which elicit a stronger response in the SC as well as larger magnitude EVRs compared to low contrast images (Kozak & Corneil, 2021; Nguyen et al., 2014; Wood et al., 2015). We predicted that we would observe a larger face-related effect in experiment 2 than experiment 1. As can be seen in the upper two plots of figure 5B, our frequentist analysis finds a significant influence of block instruction given the same visual stimuli in both the “Choose Face” and “Choose Scramble” blocks (e.g., significant effects (p< 0.01) are seen when the instructed stimulus appears on the left). Similarly, using the bayesian analysis in the bottom 2 rows of figure 5B, red banding during the EVR epoch in the upper plot indicates a consistent effect of block instruction with iterative sampling. In the bottom plot, for both comparisons the BF peaks during the EVR epoch above 100, indicating an “extreme” likelihood of an effect on block instruction on EVR expression (face left: BF=636, t=101ms; scramble left: BF=1328, t=112ms). Therefore, there was an “extreme” likelihood that block instruction resulted in a higher EMG magnitude when the participant is instructed to reach to the leftward image vs. instructed to reach to the rightward image, regardless of block instruction.

Could our results from Experiment 1 and 2 simply be due to early voluntary muscle activity influencing muscle recruitment during the EVR epoch? Two aspects of our results argue against this. First, the vast majority of RTs on target-distractor trials are > 200 ms; as with previous work we excluded trials with RTs between 120 and 150 ms to ensure that voluntary muscle activity did not influence recruitment during the EVR epoch. Second, the timecourse of the BF through time does not simply increase monotonically. Instead, for both Experiment 1 and 2, after the peak during the EVR epoch, the BF decreases to ∼1 at around 140ms following stimulus onset. It then increases again around the time of voluntary movement onset (150ms). Thus, the difference in EMG magnitude observed during the EVR epoch is distinct from that observed due to voluntary movement activity that proceeds it.

#### Group analysis: Experiments 3

Experiments 1 and 2 used highly salient repeated targets. Experiment 3 aimed to test if the face effect would extend to non-repeated, less salient targets. In any given trial of experiment 3, the targets and distractor were unpredictable to the participant, whereby any one of ten faces or ten cars could be presented as target or distractor. Unlike experiments 1 and 2, there was no significant effect of block instruction on the EVR in the frequentist analysis. Similarly, the BF did not indicate a strong likelihood of an effect of block instruction or the null hypothesis (1/10<BF<10).

#### Group analysis: Experiments 4 and 5

In experiment 2, we saw an effect of block instruction when one image was a face and the other was a scrambled image. Perhaps the EVR would reflect a similar preference for any instructed target, independent of content, as long as the target is predictable (unlike in experiment 3), or perhaps the strong effects in Experiment 2 reflect some aspect of facial affect. We therefore conducted experiments 4 and 5. In experiment 4 we used two highly salient abstract objects (+ Vs. x), and in experiment 5 we used face symbols of different affect (happy vs. Sad). No effect of block instruction was observed for experiments 4 and 5. No significant effect was found in our frequentist analysis and the BF indicates that the likelihood that block instruction had any effect on muscle activity during the EVR epoch (80-120ms) was “anecdotal” for either M_1_ or M_0_ (Fig. 5D,E; 1/10<BF<10).

### EVRs facilitate lower latency voluntary responses

The EVR has been found in previous studies to correlate with RT, consistent with the EVR leading to the development of forces that aid arm movement initiation (Contemori et al., 2021; Kearsley et al., 2022; Kozak & Corneil, 2021). To contextualize our findings, we conducted a cross-experiment analysis of the correlation coefficient between RT and EMG magnitude during the EVR epoch (80–120ms). We found that there was a significant negative correlation between RT and EVR magnitude when the instructed target was on the left (p<0.001) and a significant positive correlation when the instructed target was on the right (p<0.001), using a two-tailed one-sample t-test (Fig. 6).

**Figure 6.**
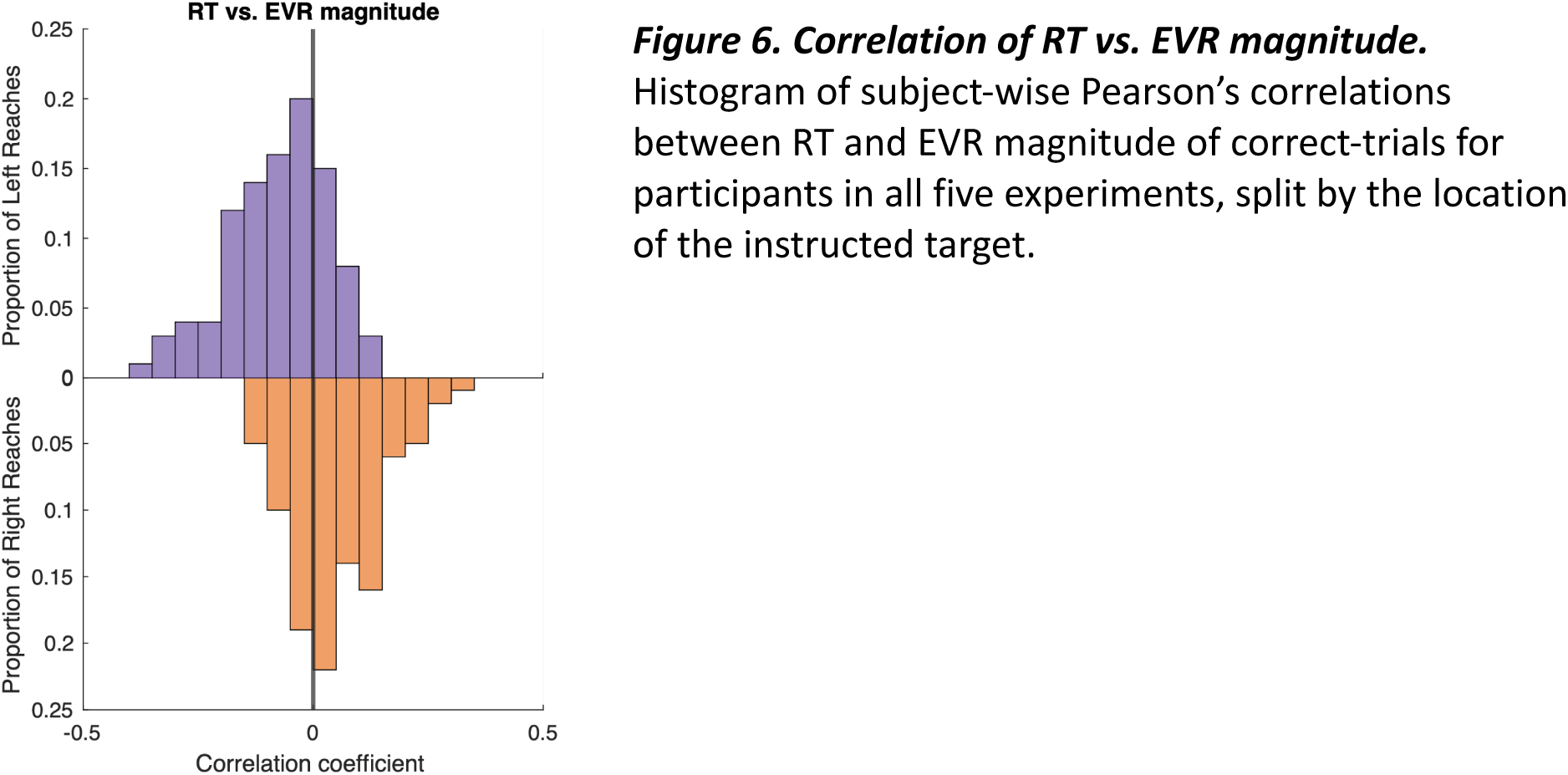
Correlation of RT vs. EVR magnitude. Histogram of subject-wise Pearson’s correlations between RT and EVR magnitude of correct-trials for participants in all five experiments, split by the location of the instructed target.

### Alternative Group Analysis

While we did observe an effect due to block instruction in experiments 1 and 2 with our first analysis, we were concerned that our results trial exclusion criteria inadvertently biased the results. This possibility is unlikely, since the analysis did not yield a similar effect in experiments 3-5, despite similar trail exclusion rates. However, to confirm the validity of our finding, we repeated all our analyses after adopting less conservative trial exclusion criteria, excluding trials where the participant moved too early (before 150ms) or where they completed the reach towards the wrong target (∼3% trial exclusion rate), but including trials where the participants moved part way toward the distractor before correctly reaching toward the target (these trials constituted 22% of all trials).

Despite the inclusion of these trials, we observe similar trends in our analysis. For experiments 1 and 2, the frequentist analysis indicates a significant effect of block instruction when the face is on the left and the scrambled image is on the right (Fig. 7A,B). Our Bayesian analysis indicates that there is a “strong” likelihood of an effect of block instruction during the EVR epoch when the face was on the left in experiment 1 (face left: BF=29,t=113ms; scramble left=1.0, t=98ms) and a “very strong” and “moderate” likelihood of an effect of block instruction in both comparisons in experiment 2 (face left: BF=38, t=101ms; scramble left: BF=6, t=108ms). As with our initial analysis, the alternative analysis yielded null findings in experiments 3-5 (Fig. 7C-E).

**Figure 7.**
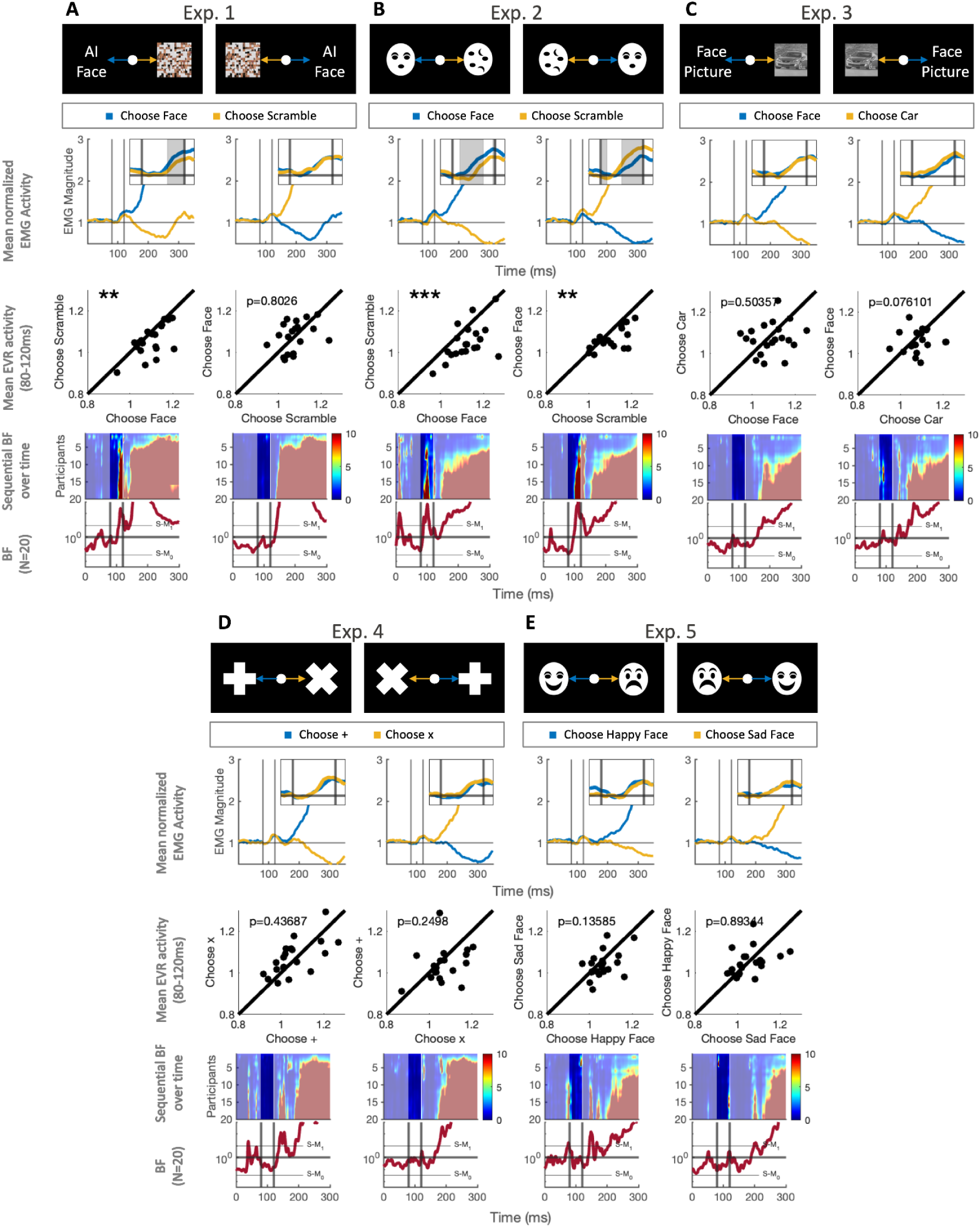
Alternative Group Analyses for Experiments 1-5. Group analysis as in figure 5, including trials where the participants moved part-way towards the distractor.

## DISCUSSION

Our hypothesis is that the EVR is evoked by the tecto-reticulo-spinal pathway. Our prediction of a face preference for the EVR was inspired by findings of increased express saccades to faces (Crouzet et al., 2010; Crouzet & Thorpe, 2011) and enhanced visual responses to faces in the SC (Nguyen et al., 2014, 2016; Van Le et al., 2020; Yu et al., 2023). Across five experiments, we quantified the EVR in a task requiring participants to reach toward an instructed target in the presence of a distractor. Across blocks, instruction varied as to which stimulus was the target. We found that block instruction influenced the EVR when one target was a salient, repeated face (Fig. 5A,B), in support of our hypothesis.

### EVRs are evoked by diametrically opposed stimuli

Our experimental findings rely on the EVR being evoked on target & distractor trials. We did indeed find this, confirming recent results of EVRs evoked by paired visual stimuli separated by 180 radial degrees in a planar reaching task (Selen et al., 2023). We presume that the EVR to paired stimuli results from how the tecto-reticulo-spinal pathway integrates visually-related activity simultaneously in each SC. When stimuli are equally weighted (e.g., Experiments 3-5), an uninfluenced EVR is evoked (Fig. 5C-E). This stimulus weighting may be biased by the nature of the stimulus and block instruction.

We did not measure muscles that act as antagonists to the pectoralis muscle, hence do not know if co-contraction is occurring during the EVR interval. Our loading of the pectoralis muscle meant that the EVR could consist of an increase or decrease in recruitment. The correlations we observed (Fig. 6) shows that the EVR produced forces that impacted subsequent reaching RTs.

### Faces impact the EVR in a task-dependent fashion

Our main findings for an influence of faces on the EVR come from Experiment 1 and 2 (Fig. 5A, B), where we found larger EVRs following leftward presentation of the instructed target. Follow-up experiments demonstrated that such a preference could not be attributable to task difficulty or be observed with any instructed object, as an instructed effect on the EVR to repeatable, salient objects was not observed in Experiment 4 (Fig. 5D) despite similar RTs and error rates as Experiment 2. In Experiment 5 (Fig. 5E), our observation of no influence of instructed facial affect on the EVR resonate with recent studies questioning the “threat bias” theory (Bannerman et al., 2009, 2010; Hansen & Hansen, 1988; Niedenthal, 1990), finding that methodological controls eliminate saccadic preferences towards angry faces (Coelho et al., 2010; Thierry et al., 2007; Webb et al., 2022). This indicates that the mechanism underlying the EVR can distinguish between evolutionary relevant objects (faces) and abstract objects (scrambled images), but cannot distinguish between abstract objects nor faces of different affect.

### The importance of stimulus salience and predictability

While block instruction influenced EVR magnitude in Experiments 1 and 2, no effect was observed in Experiment 3. We speculate that these results attest to the importance of stimulus salience and predictability. Our strongest results are in Experiment 2, which used high contrast face-like stimuli; an influence of instruction produced larger EVR when the instructed stimulus appeared on the left, regardless of instruction. In Experiment 1, we observed a similar but smaller effect on EVR expression only when the face was on the left and the scrambled image was on the right. The results in Experiments 1 and 2 are consistent with studies showing that low-spatial frequency, high-contrast, and more salient stimuli evoke larger magnitude EVRs, more express saccades, and stronger visual responses in the movement-related layers of the SC (Chen et al., 2018; Marino et al., 2012; Meeter & Van der Stigchel, 2013; Webb et al., 2022).

Experiments 2 and 3 demonstrate the importance of predictability. In Exp. 2, the two images (face vs. scramble) did not change across all 8 blocks. In Exp. 3, 20 images of faces and cars were randomly presented, making the target and distractor unpredictable. In Exp. 3 we did not observe an effect of block instruction on the EVR as we did in Exp. 2. The effect of target predictability was also observed behaviorally, as reach RT was ∼75ms longer in Exp. 3 in target & distractor conditions (Fig 3B).

### Top-down and bottom-up effects on the EVR

The EVR is influenced by both bottom-up processes related to stimulus properties and top-down instructions (Contemori et al., 2023). This resembles the process of express saccade generation, wherein the magnitude of the express response depends on both the bottom-up visual response in the SC and baseline activity in the SC as established by task set (Munoz et al., 2000). Our current findings reinforce that the top-down signal modulates the bottom-up responses when faces are present; this modulation is necessary to explain results from Experiment 2, where a larger EVR is evoked when the instructed stimulus appears on the left in both the ‘Choose Face’ and ‘Choose Scramble’ blocks. We surmise that the instruction to move to the scrambled face is akin to responding in an anti-reach task (Gu et al., 2016), wherein the EVR is muted.

We presume such a top-down influence in the setting of SC activity comes from ample projections from the frontal cortex, which is involved in the implementation and selection of task sets (Cameron et al., 2015; Johnston & Everling, 2006; Postle, 2005; Wallis et al., 2001) and basal ganglia, which codes for task urgency and reward expectation (Hikosaka et al., 2000; Kawagoe et al., 1998; Thura & Cisek, 2017; Yasuda & Hikosaka, 2015). It is unlikely that these areas produce the bottom-up effect since EVR latencies are shorter than visual response and face detection latencies in the primate prefrontal cortex (Blanke et al., 1999; Heekeren et al., 2006) and basal ganglia (Kunimatsu et al., 2023).

While visual response latencies in the SC are short enough to produce the EVR (Cecala & Freedman, 2008), it is unclear if this arises from retinotectal projections to the superficial SC, as in mice (Cazemier et al., 2022), or from the retino-geniculo-striate pathway (White et al., 2017). Recent findings that SC face responses disappear after temporary lesions of the lateral geniculate nucleus (LGN; (Yu et al., 2023) are not definitive, since such lesions abolish all bottom-up visual responses in the SC (Schiller et al., 1974). The loss of SC visual responses after LGN inactivation could arise from the loss of reticulo-geniculo-striate signalling and/or an increase in contralesional SC inhibition of the ipsilesional SC as seen in the Sprague effect (Goodale, 1973; Schiller et al., 1974; Sprague, 1966).

### Evolutionary underpinning of rapid face-related orienting

Despite the physiological and behavioral correlates supporting our finding, a key question remains: what advantage does the EVR’s ability to facilitate reaching towards or away from faces confer? A possible answer lies in literature on the superior colliculus (SC) and its role in reaching and full-body movement. The SC contains reach and grasp neurons, firing strongly when the hand touches an object in their receptive field (Kutz et al., 1997; Nagy et al., 2006; Stuphorn et al., 1999, 2000; Werner, 1993; Werner, Dannenberg, et al., 1997). SC microstimulation can evoke varying degrees of full-body movement across a variety of species, including monkeys, cats, rodents, snakes, frogs, goldfish, and lampreys (Courjon et al., 2004; Cowie & Robinson, 1994; Dacey & Ulinski, 1986; Dean et al., 1986; Ewert & Arbib, 2013; Herrero et al., 1998; Saitoh et al., 2007; Schaefer, 1970; Syka & Radil-Weiss, 1971; Tehovnik & Yeomans, 1986). Activity in the primate SC and mesencephalic reticular formation correlates with the EMG signals in the upper limbs (Stuphorn et al., 1999; Werner, Hoffmann, et al., 1997). Recent studies in humans have reported that EVRs can be distributed bilaterally in reaching tasks (Kearsley et al., 2022) and lower limbs during visually guided stepping (Billen et al., 2023), consistent with the involvement of the reticulospinal system.

Our results indicate that early face detection in the SC may serve the ecological purpose of aiding rapid orienting of the gaze axis. To the best of our knowledge, the SC is the only region projecting directly to the reticular formation that can both rapidly identify a target (Nguyen et al., 2014, 2016; Van Le et al., 2020; Yu et al., 2023) and direct orienting movements (Bogadhi et al., 2021; Herman et al., 2018).

## Conclusion

We conclude that the sensorimotor pathway underlying the EVR can rapidly initiate reaching in response to faces in a task-dependent manner. This pathway does so at latencies far shorter than those associated with face detection in the cortex (Kato et al., 2004). Previous electrophysiological studies have shown a short latency response to faces in the primate SC (Nguyen et al., 2014, 2016; Yu et al., 2023). This evidence has been interpreted as support for the idea that the SC aids cortical face detection by tagging (Yu et al., 2023) or preferentially filtering (Nguyen et al., 2014, 2016) relevant spatial locations via ascending tecto-thalamo-cortical projections. Our results align with this work and link the rapid processing of faces in the SC to rapid recruitment of upper limb muscles.

## Acknowledgments

This work is supported by Discovery Grants to BDC from the Natural Sciences and Engineering Research Council of Canada (NSERC; RGPIN 311680 and 04394-2021) and an Operating Grant to BDC from the Canadian Institutes of Health Research (CIHR; MOP-93796). The equipment used in this experiment was purchased using funds from the Canadian Foundation for Innovation. Additional support came from the Canada First Research Excellence Fund (BrainsCAN). Special thanks to Drs. Aaron L. Cecala, Rebecca Kozak, Paul L. Gribble, Andrew J. Pruszynski, Jody C. Culham, Guy Wallis, Gerald E. Loeb, and Timothy J. Carroll for their guidance and mentorship.

## REFERENCES

Almeida, I., Soares, S. C., & Castelo-Branco, M. (2015). The Distinct Role of the Amygdala, Superior Colliculus and Pulvinar in Processing of Central and Peripheral Snakes. PloS One, 10(6), e0129949.

Bannerman, R. L., Milders, M., de Gelder, B., & Sahraie, A. (2009). Orienting to threat: faster localization of fearful facial expressions and body postures revealed by saccadic eye movements. Proceedings. Biological Sciences / The Royal Society, 276(1662), 1635–1641.

Bannerman, R. L., Milders, M., & Sahraie, A. (2010). Attentional bias to brief threat-related faces revealed by saccadic eye movements. Emotion, 10(5), 733–738.

Bentin, S., Allison, T., Puce, A., Perez, E., & McCarthy, G. (1996). Electrophysiological Studies of Face Perception in Humans. Journal of Cognitive Neuroscience, 8(6), 551–565.

Billen, L. S., Corneil, B. D., & Weerdesteyn, V. (2023). Evidence for an Intricate Relationship Between Express Visuomotor Responses, Postural Control and Rapid Step Initiation in the Lower Limbs. Neuroscience, 531, 60–74.

Blanke, O., Morand, S., Thut, G., Michel, C. M., Spinelli, L., Landis, T., & Seeck, M. (1999). Visual activity in the human frontal eye field. Neuroreport, 10(5), 925–930.

Bogadhi, A. R., & Hafed, Z. M. (2022). Express detection and discrimination of visual objects by primate superior colliculus neurons. In bioRxiv. 10.1101/2022.02.08.479583

Bogadhi, A. R., Katz, L. N., Bollimunta, A., Leopold, D. A., & Krauzlis, R. J. (2021). Midbrain activity shapes high-level visual properties in the primate temporal cortex. Neuron, 109(4), 690–699.e5.

Cameron, I. G. M., Riddle, J. M., & D’Esposito, M. (2015). Dissociable Roles of Dorsolateral Prefrontal Cortex and Frontal Eye Fields During Saccadic Eye Movements. Frontiers in Human Neuroscience, 9, 613.

Cazemier, J. L., Tran, T. K. L., Hsu, A. T. Y., Husić, M., Kirchberger, L., Self, M. W., Roelfsema, P. R., & Heimel, J. A. (2022). Involvement of superior colliculus in complex figure detection of mice. In bioRxiv. 10.1101/2022.09.25.509365

Cecala, A. L., & Freedman, E. G. (2008). Amplitude changes in response to target displacements during human eye-head movements. Vision Research, 48(2), 149–166.

Celeghin, A., Bagnis, A., Diano, M., Méndez, C. A., Costa, T., & Tamietto, M. (2019). Functional neuroanatomy of blindsight revealed by activation likelihood estimation meta-analysis. Neuropsychologia, 128, 109–118.

Chen, C.-Y., Sonnenberg, L., Weller, S., Witschel, T., & Hafed, Z. M. (2018). Spatial frequency sensitivity in macaque midbrain. Nature Communications, 9(1), 2852.

Coelho, C. M., Cloete, S., & Wallis, G. (2010). The face-in-the-crowd effect: when angry faces are just cross(es). Journal of Vision, 10(1), 7.1–14.

Collins, J. A., & Olson, I. R. (2014). Beyond the FFA: The role of the ventral anterior temporal lobes in face processing. Neuropsychologia, 61, 65–79.

Contemori, S., Loeb, G. E., Corneil, B. D., Wallis, G., & Carroll, T. J. (2021). Trial-by-trial modulation of express visuomotor responses induced by symbolic or barely detectable cues. Journal of Neurophysiology, 126(5), 1507–1523.

Contemori, S., Loeb, G. E., Corneil, B. D., Wallis, G., & Carroll, T. J. (2023). Express Visuomotor Responses Reflect Knowledge of Both Target Locations and Contextual Rules during Reaches of Different Amplitudes. The Journal of Neuroscience: The Official Journal of the Society for Neuroscience, 43(42), 7041–7055.

Corneil, B. D., & Munoz, D. P. (2014). Overt responses during covert orienting. Neuron, 82(6), 1230–1243.

Courjon, J.-H., Olivier, E., & Pélisson, D. (2004). Direct evidence for the contribution of the superior colliculus in the control of visually guided reaching movements in the cat. The Journal of Physiology, 556(Pt 3), 675–681.

Cowie, R. J., & Robinson, D. L. (1994). Subcortical contributions to head movements in macaques. I. Contrasting effects of electrical stimulation of a medial pontomedullary region and the superior colliculus. Journal of Neurophysiology, 72(6), 2648–2664.

Crouzet, S. M., Kirchner, H., & Thorpe, S. J. (2010). Fast saccades toward faces: face detection in just 100 ms. Journal of Vision, 10(4), 16.1–17.

Crouzet, S. M., & Thorpe, S. J. (2011). Low-level cues and ultra-fast face detection. Frontiers in Psychology, 2, 342.

Dacey, D. M., & Ulinski, P. S. (1986). Optic tectum of the eastern garter snake, Thamnophis sirtalis. V. Morphology of brainstem afferents and general discussion. The Journal of Comparative Neurology, 245(4), 423–453.

Dean, P., Redgrave, P., Sahibzada, N., & Tsuji, K. (1986). Head and body movements produced by electrical stimulation of superior colliculus in rats: effects of interruption of crossed tectoreticulospinal pathway. Neuroscience, 19(2), 367–380.

Decramer, T., Premereur, E., Zhu, Q., Van Paesschen, W., van Loon, J., Vanduffel, W., Taubert, J., Janssen, P., & Theys, T. (2021). Single-Unit Recordings Reveal the Selectivity of a Human Face Area. The Journal of Neuroscience: The Official Journal of the Society for Neuroscience, 41(45), 9340–9349.

Ewert, J. P., & Arbib, M. A. (2013). Visuomotor Coordination: Amphibians, Comparisons, Models, and Robots. Springer Science & Business Media.

Gandhi, N. J., & Katnani, H. A. (2011). Motor functions of the superior colliculus. Annual Review of Neuroscience, 34, 205–231.

Goodale, M. A. (1973). Cortico-tectal and intertectal modulation of visual responses in the rat’s superior colliculus. Experimental Brain Research. Experimentelle Hirnforschung. Experimentation Cerebrale, 17(1), 75–86.

Gu, C., Wood, D. K., Gribble, P. L., & Corneil, B. D. (2016). A Trial-by-Trial Window into Sensorimotor Transformations in the Human Motor Periphery. The Journal of Neuroscience: The Official Journal of the Society for Neuroscience, 36(31), 8273–8282.

Hansen, C. H., & Hansen, R. D. (1988). Finding the face in the crowd: an anger superiority effect. Journal of Personality and Social Psychology, 54(6), 917–924.

Heekeren, H. R., Marrett, S., Ruff, D. A., Bandettini, P. A., & Ungerleider, L. G. (2006). Involvement of human left dorsolateral prefrontal cortex in perceptual decision making is independent of response modality. Proceedings of the National Academy of Sciences of the United States of America, 103(26), 10023–10028.

Herman, J. P., Katz, L. N., & Krauzlis, R. J. (2018). Midbrain activity can explain perceptual decisions during an attention task. Nature Neuroscience, 21(12), 1651–1655.

Herrero, L., Rodríguez, F., Salas, C., & Torres, B. (1998). Tail and eye movements evoked by electrical microstimulation of the optic tectum in goldfish. Experimental Brain Research. Experimentelle Hirnforschung. Experimentation Cerebrale, 120(3), 291–305.

Hikosaka, O., Takikawa, Y., & Kawagoe, R. (2000). Role of the basal ganglia in the control of purposive saccadic eye movements. Physiological Reviews, 80(3), 953–978.

Jeffreys, H. (1998). The Theory of Probability. OUP Oxford.

Johnston, K., & Everling, S. (2006). Monkey dorsolateral prefrontal cortex sends task-selective signals directly to the superior colliculus. The Journal of Neuroscience: The Official Journal of the Society for Neuroscience, 26(48), 12471–12478.

Kato, Y., Muramatsu, T., Kato, M., Shintani, M., Yoshino, F., Shimono, M., & Ishikawa, T. (2004). An earlier component of face perception detected by seeing-as-face task. Neuroreport, 15(2), 225–229.

Kawagoe, R., Takikawa, Y., & Hikosaka, O. (1998). Expectation of reward modulates cognitive signals in the basal ganglia. Nature Neuroscience, 1(5), 411–416.

Kearsley, S. L., Cecala, A. L., Kozak, R. A., & Corneil, B. D. (2022). Express arm responses appear bilaterally on upper-limb muscles in an arm choice reaching task. Journal of Neurophysiology, 127(4), 969–983.

Kozak, R. A., Cecala, A. L., & Corneil, B. D. (2020). An Emerging Target Paradigm to Evoke Fast Visuomotor Responses on Human Upper Limb Muscles. Journal of Visualized Experiments: JoVE, 162. 10.3791/61428

Kozak, R. A., & Corneil, B. D. (2021). High-contrast, moving targets in an emerging target paradigm promote fast visuomotor responses during visually guided reaching. Journal of Neurophysiology, 126(1), 68–81.

Kozak, R. A., Kreyenmeier, P., Gu, C., Johnston, K., & Corneil, B. D. (2019). Stimulus-Locked Responses on Human Upper Limb Muscles and Corrective Reaches Are Preferentially Evoked by Low Spatial Frequencies. eNeuro, 6(5). 10.1523/ENEURO.0301-19.2019

Krekelberg, B. (2022). *BayesFactor: Release 2022* (v2.3.0). Zenodo. 10.5281/ZENODO.7006300

Kunimatsu, J., Amita, H., & Hikosaka, O. (2023). Neuronal mechanism of the encoding of socially familiar faces in the striatum tail. bioRxiv : The Preprint Server for Biology. 10.1101/2023.05.10.540108

Kutz, D. F., Dannenberg, S., Werner, W., & Hoffmann, K. P. (1997). Population coding of arm-movement-related neurons in and below the superior colliculus of Macaca mulatta. Biological Cybernetics, 76(5), 331–337.

Marino, R. A., Levy, R., Boehnke, S., White, B. J., Itti, L., & Munoz, D. P. (2012). Linking visual response properties in the superior colliculus to saccade behavior. The European Journal of Neuroscience, 35(11), 1738–1752.

Martin, J. G., Davis, C. E., Riesenhuber, M., & Thorpe, S. J. (2018). Zapping 500 faces in less than 100 seconds: Evidence for extremely fast and sustained continuous visual search. Scientific Reports, 8(1), 12482.

Meeter, M., & Van der Stigchel, S. (2013). Visual priming through a boost of the target signal: evidence from saccadic landing positions. Attention, Perception & Psychophysics, 75(7), 1336–1341.

Munoz, D. P., Dorris, M. C., Paré, M., & Everling, S. (2000). On your mark, get set: brainstem circuitry underlying saccadic initiation. Canadian Journal of Physiology and Pharmacology, 78(11), 934–944.

Nagy, A., Kruse, W., Rottmann, S., Dannenberg, S., & Hoffmann, K.-P. (2006). Somatosensory-motor neuronal activity in the superior colliculus of the primate. Neuron, 52(3), 525–534.

Nguyen, M. N., Hori, E., Matsumoto, J., Tran, A. H., Ono, T., & Nishijo, H. (2013). Neuronal responses to face-like stimuli in the monkey pulvinar. The European Journal of Neuroscience, 37(1), 35–51.

Nguyen, M. N., Matsumoto, J., Hori, E., Maior, R. S., Tomaz, C., Tran, A. H., Ono, T., & Nishijo, H. (2014). Neuronal responses to face-like and facial stimuli in the monkey superior colliculus. Frontiers in Behavioral Neuroscience, 8, 85.

Nguyen, M. N., Nishimaru, H., Matsumoto, J., Van Le, Q., Hori, E., Maior, R. S., Tomaz, C., Ono, T., & Nishijo, H. (2016). Population Coding of Facial Information in the Monkey Superior Colliculus and Pulvinar. Frontiers in Neuroscience, 10, 583.

Niedenthal, P. M. (1990). Implicit perception of affective information. Journal of Experimental Social Psychology, 26(6), 505–527.

Postle, B. R. (2005). Delay-period activity in the prefrontal cortex: one function is sensory gating. Journal of Cognitive Neuroscience, 17(11), 1679–1690.

Pruszynski, J. A., King, G. L., Boisse, L., Scott, S. H., Flanagan, J. R., & Munoz, D. P. (2010). Stimulus-locked responses on human arm muscles reveal a rapid neural pathway linking visual input to arm motor output. The European Journal of Neuroscience, 32(6), 1049–1057.

Saitoh, K., Ménard, A., & Grillner, S. (2007). Tectal control of locomotion, steering, and eye movements in lamprey. Journal of Neurophysiology, 97(4), 3093–3108.

Salvia, E., Grosbras, M. H., Harvey, M., & Nazarian, B. (2020). Social perception drives eye-movement related brain activity: Evidence from pro- and anti-saccades to faces. Neuropsychologia, 139, 107360.

Schaefer, K. P. (1970). Unit analysis and electrical stimulation in the optic tectum of rabbits and cats. Brain, Behavior and Evolution, 3(1), 222–240.

Schiller, P. H., Stryker, M., Cynader, M., & Berman, N. (1974). Response characteristics of single cells in the monkey superior colliculus following ablation or cooling of visual cortex. Journal of Neurophysiology, 37(1), 181–194.

Selen, L. P. J., Corneil, B. D., & Medendorp, W. P. (2023). Single-Trial Dynamics of Competing Reach Plans in the Human Motor Periphery. The Journal of Neuroscience: The Official Journal of the Society for Neuroscience, 43(15), 2782–2793.

Sprague, J. M. (1966). Interaction of cortex and superior colliculus in mediation of visually guided behavior in the cat. Science, 153(3743), 1544–1547.

Stuphorn, V., Bauswein, E., & Hoffmann, K. P. (2000). Neurons in the primate superior colliculus coding for arm movements in gaze-related coordinates. Journal of Neurophysiology, 83(3), 1283–1299.

Stuphorn, V., Hoffmann, K. P., & Miller, L. E. (1999). Correlation of primate superior colliculus and reticular formation discharge with proximal limb muscle activity. Journal of Neurophysiology, 81(4), 1978–1982.

Syka, J., & Radil-Weiss, T. (1971). Electrical stimulation of the tectum in freely moving cats. Brain Research, 28(3), 567–572.

Tehovnik, E. J., & Yeomans, J. S. (1986). Two converging brainstem pathways mediating circling behavior. Brain Research, 385(2), 329–342.

Thierry, G., Martin, C. D., Downing, P., & Pegna, A. J. (2007). Controlling for interstimulus perceptual variance abolishes N170 face selectivity. Nature Neuroscience, 10(4), 505–511.

Thura, D., & Cisek, P. (2017). The Basal Ganglia Do Not Select Reach Targets but Control the Urgency of Commitment. Neuron, 95(5), 1160–1170.e5.

Van Le, Q., Van Le, Q., Nishimaru, H., Matsumoto, J., Takamura, Y., Hori, E., Maior, R. S., Tomaz, C., Ono, T., & Nishijo, H. (2020). A Prototypical Template for Rapid Face Detection Is Embedded in the Monkey Superior Colliculus. Frontiers in Systems Neuroscience, 14, 507700.

VanRullen, R., & Thorpe, S. J. (2001). The time course of visual processing: from early perception to decision-making. Journal of Cognitive Neuroscience, 13(4), 454–461.

Wallis, J. D., Anderson, K. C., & Miller, E. K. (2001). Single neurons in prefrontal cortex encode abstract rules. Nature, 411(6840), 953–956.

Webb, A. L. M., Asher, J. M., & Hibbard, P. B. (2022). Saccadic eye movements are deployed faster for salient facial stimuli, but are relatively indifferent to their emotional content. Vision Research, 198, 108054.

Werner, W. (1993). Neurons in the primate superior colliculus are active before and during arm movements to visual targets. The European Journal of Neuroscience, 5(4), 335–340.

Werner, W., Dannenberg, S., & Hoffmann, K. P. (1997). Arm-movement-related neurons in the primate superior colliculus and underlying reticular formation: comparison of neuronal activity with EMGs of muscles of the shoulder, arm and trunk during reaching. Experimental Brain Research. Experimentelle Hirnforschung. Experimentation Cerebrale, 115(2), 191–205.

Werner, W., Hoffmann, K. P., & Dannenberg, S. (1997). Anatomical distribution of arm-movement-related neurons in the primate superior colliculus and underlying reticular formation in comparison with visual and saccadic cells. Experimental Brain Research. Experimentelle Hirnforschung. Experimentation Cerebrale, 115(2), 206–216.

White, B. J., Kan, J. Y., Levy, R., Itti, L., & Munoz, D. P. (2017). Superior colliculus encodes visual saliency before the primary visual cortex. Proceedings of the National Academy of Sciences of the United States of America, 114(35), 9451–9456.

Wood, D. K., Gu, C., Corneil, B. D., Gribble, P. L., & Goodale, M. A. (2015). Transient visual responses reset the phase of low-frequency oscillations in the skeletomotor periphery. The European Journal of Neuroscience, 42(3), 1919–1932.

Yasuda, M., & Hikosaka, O. (2015). Functional territories in primate substantia nigra pars reticulata separately signaling stable and flexible values. Journal of Neurophysiology, 113(6), 1681–1696.

Yu, G., Katz, L. N., Quaia, C., Messinger, A., & Krauzlis, R. J. (2023). Short-latency preference for faces in the primate superior colliculus. In bioRxiv. 10.1101/2023.09.06.556401

Zellner, A., & Siow, A. (1979). On Posterior Odds Ratios for Sharp Null Hypothesis and One-sided Alternatives.

